# Targeted Perturb-seq Reveals EGR1 and FOS as Key Regulators of the Transcriptional RAF-MAPK Response

**DOI:** 10.1101/2024.01.13.575500

**Authors:** Ghanem El Kassem, Anja Sieber, Bertram Klinger, Florian Uhlitz, David Steinbrecht, Mirjam van Bentum, Jasmine Hillmer, Jennifer von Schlichting, Reinhold Schäfer, Nils Blüthgen, Michael Boettcher

## Abstract

The MAPK pathway is an important cellular signaling cascade whose dysregulation causes a variety of diseases. While the upstream regulators of this cascade have been extensively characterized, the understanding of how its activation translates into different transcriptional responses remains poorly understood. This study attempts to fill this knowledge gap by using targeted Perturb-seq against 22 transcription factors in an inducible model system for RAF-MAPK signaling. A topology-based modeling approach is applied to the obtained data to construct a directional interaction network. By removing coherent feed-forward loops and integrating the expression kinetics of transcription factors, a parsimonious network structure is derived that distinguishes direct from indirect interactions between the investigated transcription factors and their targets. In particular, EGR1 and FOS are found to act as orthogonal upstream regulators of the RAF-MAPK response. The results presented here provide valuable insights into the organization of the transcriptional network downstream of RAF-MAPK signaling and thus provide a basis for a better understanding of this complex process.

## Introduction

The Mitogen-Activated Protein Kinase (MAPK) signaling cascade is a fundamental and evolutionarily conserved pathway crucial for the regulation of key cellular processes, including proliferation, differentiation, and survival (Cargnello & Roux, 2011). Triggered by external stimuli such as growth factors, hormones, and stress, this pathway transmits these signals from the cell surface into the nucleus. Upon activation, MAP kinases relocate into the nucleus and phosphorylate transcription factors, thereby activating specific transcriptional programs. The adaptability of MAPK signaling allows cells to effectively integrate a broad range of external signals into cellular responses. Dysregulation of the MAPK signaling pathway is implicated in various diseases, including cancer, neurodegenerative disorders, and inflammatory conditions (Kim & Choi, 2010; Dhillon *et al*, 2007). Therefore, MAPK signaling is an attractive target for therapeutic intervention, aimed at restoring normal signaling patterns and mitigating the effects of aberrant signaling on disease progression (Burotto *et al*, 2014; Bahar *et al*, 2023).

The MAPK pathway has served as a paradigm for studying signal-induced transcriptional programs (Schulze *et al*, 2001). Early transcriptome studies after stimulation of the pathway led to the concepts of immediate-early and delayed primary genes that are activated by pre-existing transcription factors (Tullai *et al*, 2007), and share regulatory motifs in their promoters (Tullai *et al*, 2004; Jürchott *et al*, 2010). Many immediate-early genes are transcription factors, which then induce secondary targets. Interestingly, delayed primary genes often encode negative feedback regulators (Amit *et al*, 2007; Avraham & Yarden, 2011; Legewie *et al*, 2008), and the timing of primary response genes is strongly determined by mRNA half-lives (Uhlitz *et al*, 2017). While the kinetics of the transcriptome response to MAPK activation has been characterized in quantitative depth (Tullai *et al*, 2007; Amit *et al*, 2007; Uhlitz *et al*, 2017; Legewie *et al*, 2008), the wiring of how immediate-early transcription factors induce secondary response genes and the understanding of their interaction remain cryptic.

Reverse engineering of the transcriptional networks downstream of MAPK signaling represents a promising approach to better understand how physiological and pathological MAPK signals are integrated into a cellular response. We have previously used systematic perturbation data and reverse engineering to elucidate a small seven node transcription factor network downstream of RAS/MAPK signaling that controls transformation and different aspects of cell growth (Stelniec-Klotz *et al*, 2012). Expansion of such limited approaches to larger networks has been hampered by technical challenges for a long time. However, newly developed methods that combine CRISPR-based genetic perturbation techniques with single-cell RNA-Seq readout, such as Perturb-seq (Adamson *et al*, 2016; Dixit *et al*, 2016), now enable simultaneous functional genetic perturbation and investigation of the resulting transcriptional response at scale.

In this study, we employed a targeted Perturb-seq (TAP-seq) approach (Schraivogel *et al*, 2020) to explore the transcriptional outcomes associated with the knockout of 22 transcription factors induced by the MAPK signaling kinase RAF1. Among these factors, EGR1 and FOS emerged as the most upstream regulators of the transcriptional RAF-MAPK response. Interestingly, these two transcription factors frequently co-regulate overlapping target gene sets in an orthogonal manner. Collectively, our findings underscore the efficacy of targeted Perturb-seq in reconstructing the topology of the transcriptional networks that mediate the response to MAPK activation.

## Results

### Identification of transcription factors up-regulated by RAF1-induction

To identify transcripts that are up-regulated by RAF-MAPK activation, we used a previously established HEK293 cell line, termed HEK293ΔRAF1:ER, in which a tamoxifen-inducible RAF1-CR3 kinase domain was introduced (Samuels *et al*, 1993). Consequently, RAF1 activity can be precisely regulated, allowing the identification of RAF1-MAPK response genes (Fig. 1A). In contrast to cell culture systems stimulated by growth factors, this model of RAF-MAPK signaling is activated independently of the upstream G-protein RAS, thus minimizing pathway divergence and feedback mechanisms. We induced RAF activity with 4- hydroxytamoxifen (4OHT) over periods of 0.5 to 8h and monitored the changes in the transcriptome of the cells via bulk RNA-sequencing (RNA-Seq). Over the full time series, we detected a total of 1,142 significantly up-regulated genes (Fig.1B). From this dataset, 22 transcription factors (TFs) that were up-regulated at different time points after RAF1 induction were selected for further analysis. The basal expression levels of identified candidate TFs varied over several orders of magnitude and their level of up-regulation upon RAF1 induction was independent of their basal expression level (Fig.1C).

**Figure 1.**
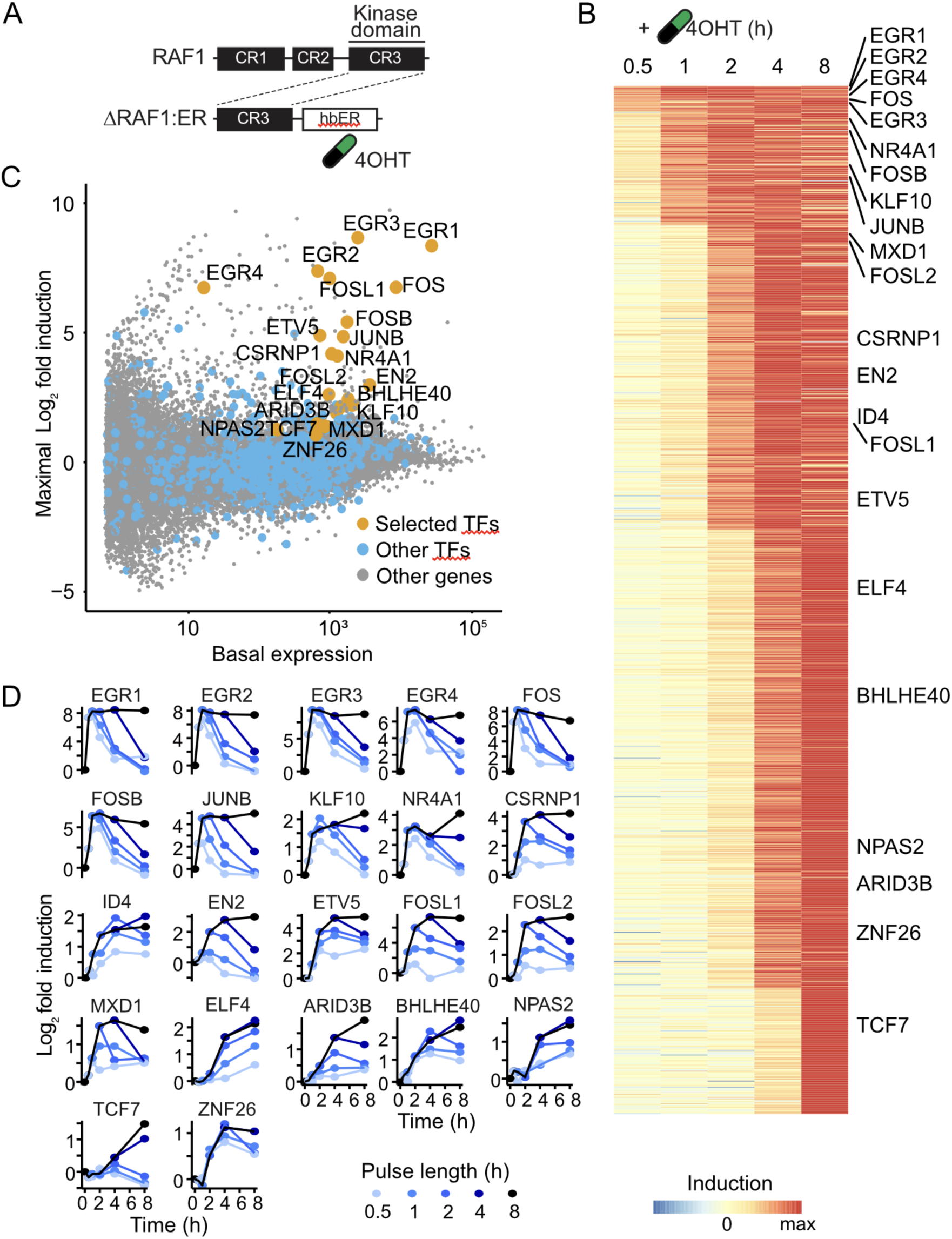
Transcriptional profiling of HEK293ΔRAF1:ER cells following RAF1 induction. **(A)** Schematic structure of the 4-hydroxytamoxifen-(4OHT)-inducible RAF1-CR3 kinase domain. **(B)** Maximum normalized log_2_ fold changes from bulk RNAseq analysis of significantly up-regulated genes (FDR<1%) after 0.5h, 1h, 2h, 4h and 8h. The 22 selected candidate TFs are indicated by their gene symbols. **(C)** The maximal log_2_ fold expression change of the selected candidate TFs is plotted against their expression levels in non-induced HEK293ΔRAF1:ER cells. **(D)** The time-resolved log_2_ fold expression changes of candidate TFs are shown after the indicated pulse lengths of 4OHT-mediated RAF1 induction.

Pulsed induction of RAF1 (Fig. 1D) further allowed the categorization of the candidate TFs into three distinct response classes. The first class comprises classical immediate early genes that are rapidly induced upon signal induction and whose mRNAs are rather short lived, resulting in rapid decay after the pulses ended. Examples of these transcripts are the rapidly induced EGR gene family members, FOS, FOSB, and JUNB, which increased to maximum mRNA levels right after induction and decreased quickly after 4OHT removal. The second class comprises rather rapidly induced, long-lived transcripts that are induced within the first 1-2h and remain high even after the pulse has ended. These include, for instance, the transcripts of FOSL1 and FOSL2. A third class of transcription factors is only induced with delay. For several of those transcripts, the response is limited to long pulses of induction. For instance, TCF7 is only induced after 4h and reaches a maximum response after 8h of induction. We furthermore calculated the time when the transcripts reach half-maximal expression by interpolating between the measured time points. We found that the selected transcripts cover half-maximal induction times between 20 min and 5h (Table 1).

**Table 1.** Selected candidate genes and their classification based on their response time to RAF1 induction (Uhlitz *et al*, 2017). IEG = immediate-early genes, ILG = immediate-late genes, DEG = delayed-early genes.

| Gene Symbol | Gene name | Class (Uhlitz *et al*, 2017) | Half-maximal induction (h:min) |
| --- | --- | --- | --- |
| EGR1 | Early Growth Response 1 | IEG | 0:17 |
| FOS | Fos Proto-Oncogene, | IEG | 0:19 |
| EGR2 | Early Growth Response 2 | IEG | 0:22 |
| EGR3 | Early Growth Response 3 | IEG | 0:25 |
| EGR4 | Early Growth Response 4 | ILG | 0:29 |
| JUNB | JunB Proto-Oncogene | IEG | 0:33 |
| FOSB | FosB Proto-Oncogene | IEG | 0:37 |
| NR4A1 | Nuclear Receptor Subfamily 4 Group A Member 1 | ILG | 0:47 |
| KLF10 | KLF Transcription Factor 10 | DEG | 0:49 |
| ID4 | Inhibitor Of DNA Binding 4 | DEG | 1:05 |
| MXD1 | MAX Dimerization Protein 1 | DEG | 1:13 |
| FOSL1 | FOS Like 1 | ILG | 1:17 |
| CSRNP1 | Cysteine And Serine Rich Nuclear Protein 1 | SRG | 1:19 |
| FOSL2 | FOS Like 2 | ILG | 1:23 |
| ETV5 | ETS Variant Transcription Factor 5 | IEG | 1:31 |
| EN2 | Engrailed Homeobox 2 | SRG | 1:37 |
| ZNF26 | Zinc Finger Protein 26 | DEG | 2:09 |
| BHLHE40 | Basic Helix-Loop-Helix Family Member E40 | DEG | 2:15 |
| ARID3B | AT-Rich Interaction Domain 3B | SRG | 3:03 |
| ELF4 | E74 Like ETS Transcription Factor 4 | SRG | 3:10 |
| NPAS2 | Neuronal PAS Domain Protein 2 | SRG | 3:16 |
| TCF7 | Transcription Factor 7 | SRG | 5:08 |

Thus the selected transcription factors cover different dynamic aspects of the RAF1-induced transcriptional response, which include transcripts previously characterized as the immediate-early (IEG), immediate-late (ILG), delayed-early (DEG), and secondary response genes (SRG) (Uhlitz *et al*, 2017). The classes and half-maximal induction times of each of the 22 candidate TFs are shown in Table 1. While their induction kinetics provide an indication of whether the respective TF candidate is required early or late in the RAF response, they do not reveal whether or how the induced TFs are functionally involved in the regulation of transcriptional networks downstream of RAF-MAPK. To answer this question, we next performed a series of Perturb-seq screens.

### Perturb-seq screens for the functional characterization of TFs up-regulated by RAF1- induction

To identify the transcriptional targets of the selected 22 candidate TFs, we performed pooled CRISPR/Cas9 screens with single cell RNA-Seq read-out, via direct-capture Perturb-seq (Replogle *et al*, 2020), as well as pooled CRISPR screens with proliferation read-out (Fig. 2A). For that purpose, we designed and cloned a pooled sgRNA library, targeting the 22 candidate TFs with 4 sgRNAs each, 10 non-target control and 10 safe cutter sgRNAs. The latter cut in gene-free regions of the genome and serve as controls for potential effects of Cas9-induced DNA double strand breaks. In addition, the library contained 4 sgRNAs against the transgene containing the RAF1-CR3 kinase domain as a positive control. We confirmed that the sgRNA library was cloned with an exceptionally narrow distribution. More than 95% of the normalized sgRNA sequence read counts fell within one order of magnitude (Fig. 2B). This narrow sgRNA library distribution is of particular importance for Perturb-seq screens, in order to obtain similar numbers of analyzable cells per sgRNA. After lentiviral packaging of the sgRNA library, the HEK293ΔRAF1:ER cells were transduced at low multiplicity of infection (MOI = 0.2) and cultured for 10 days to allow efficient perturbation of the candidate TFs (Fig. S1). Following the 10-day editing period, RAF1 was induced by 4OHT for 6h, 12h, and 18h, respectively, to capture time-resolved transcription profile changes from each of the 22 perturbed candidate TFs. The cells were then run through the 10x Genomics 3’ scRNA-Seq pipeline with “Feature Barcode technology” for the simultaneous detection of the transcriptome and the sgRNA that was expressed in each single cell. All experiments were performed in duplicate.

**Figure 2.**
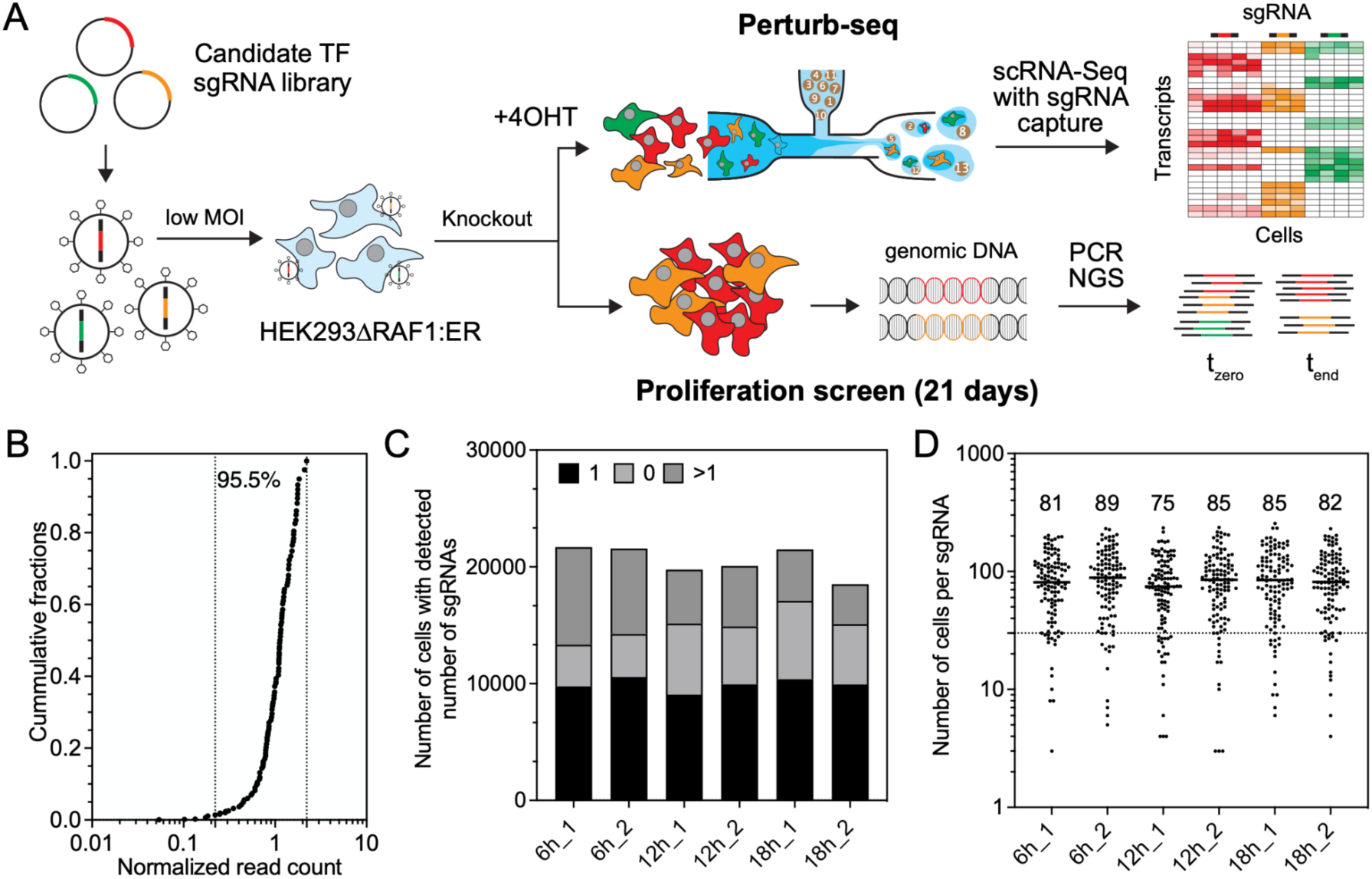
Pooled CRISPR screens in HEK293ΔRAF1:ER cells. **(A)** Schematic of Perturb-seq screens (top panel) and proliferation CRISPR screens (bottom panel). **(B)** Distribution of the pooled sgRNA library used for all Perturb-seq and proliferation screens. **(C)** Total number of recovered cells and number of cells with 0, 1, and >1 sgRNAs detected in the respective Perturb-seq samples. **(D)** Median number and distribution of cells per sgRNA detected in the respective Perturb-seq samples. Dashed line at 30 cells per sgRNA indicates the cut-off cell number per sgRNA that was used for further analysis.

As expected, the low MOI used for transduction resulted in the detection of a single sgRNA in the majority of captured cells (Fig.2C). The detection of more than one sgRNA per single cell barcode, is most likely explained by multiple cells being captured together with a 10x bead in the same emulsion droplet, a consequence of the intentionally high number of cells loaded per reaction (Fig.2C). Cells with >1 or 0 detectable sgRNA were omitted from further analysis, resulting in approximately 10,000 analyzable cells per sample with exactly one sgRNA (Fig. 2C), which was detected with a median of 26-84 UMI counts in the respective Perturb-seq samples (Fig. S2). As expected from the narrow distribution of the sgRNA library (Fig. 2B), the distribution of analyzable cells per sgRNAs was also very narrow across all six Perturb-seq samples, with median numbers ranging from 75 to 89 analyzable cells per sgRNA (Fig. 2D). Taken together, these results demonstrate the high technical quality of the generated Perturb-seq data, which formed the basis for all further analyses.

### Proliferation CRISPR screens of candidate TFs in HEK293ΔRAF1:ER cells

We first asked if TF knockouts have strong effects on cell growth, using a CRISPR/Cas9 screen with proliferation read-out (Fig. 2A, S3), both with induced RAF activity and without. We found that most TFs show no significant depletion or enrichment compared to non-targeting controls. The FOS and EGR2 guides are mildly but significantly depleted both with induced RAF activity and without. This confirms their known role in cell proliferation. Interestingly, the knockout of the ΔRAF1:ER transgene is strongly enriched, but only when the transgene is activated. This is consistent with the previous report that prolonged activation of RAF-MAPK signaling in HEK293ΔRAF1:ER cells results in apoptosis (Cagnol *et al*, 2006). Overall, this screen confirms that there is no strong depletion of guides targeting the transcription factors, allowing to perform perturb-seq experiments without losing the representation of the guide library.

### Quality control of Perturb-seq screen data

Integration of UMAPs from Perturb-seq experiments at 6h, 12h and 18h post RAF1 induction revealed that cells from the 12h and 18h time points clustered separately from those of the 6h samples (Fig. 3A). This observation suggests that major transcriptional changes happen between 6h and 12h of RAF1 induction, emphasizing the temporal dynamics of the cellular response. A uniform distribution of cells across different cell cycle phases G1, S, and G2M within the UMAP clusters indicated that cell cycle heterogeneity did not contribute to observed variations (Fig. 3B). To assess the transcriptional consequences of RAF1- induction, RAF1-transgene knockout cells were compared to safe cutter control cells. The RAF1-knockout cells formed a distinct cluster that was clearly separated from the safe cutter control cells (Fig. 3C). This separation confirms the expected absence of a transcriptional response to RAF1 induction in the RAF1 knockout cells, suggesting that the observed transcriptional changes in the safe cutter control cells are indeed due to the specific activation of the RAF1 transgene by 4OHT. Pseudo-bulk analysis of the fold changes of known RAF-MAPK targets unveiled a normally distributed peak around 0 for the safe cutter controls, while the RAF1-knockout cells showed a clear shift towards negative log_2_ fold changes (Fig. 3D). These findings illustrate that in comparison to non-target control cells, the safe cutter cells showed no transcriptional changes, suggesting no global impact of Cas9- induced DNA double strand breaks on the Perturb-seq results. Further, the clear shift of the RAF1-knockout peak towards negative log_2_ fold changes confirmed the inability of those cells to respond to 4OHT with the activation of RAF1-MAPK induced genes. Examination of the transcriptional changes of individual sgRNAs against RAF1 confirmed that all three sgRNAs were effective (the fourth RAF1 sgRNA was represented in less than 30 cells and hence was omitted from analysis), with values ranging from 12%-37% significantly deregulated genes from the total RAF1-induced genes identified from the bulk RNAseq data (Fig. 3E). Overall, these results confirm that the Perturb-seq screens were successful, as all positive and negative controls behaved as expected.

**Figure 3.**
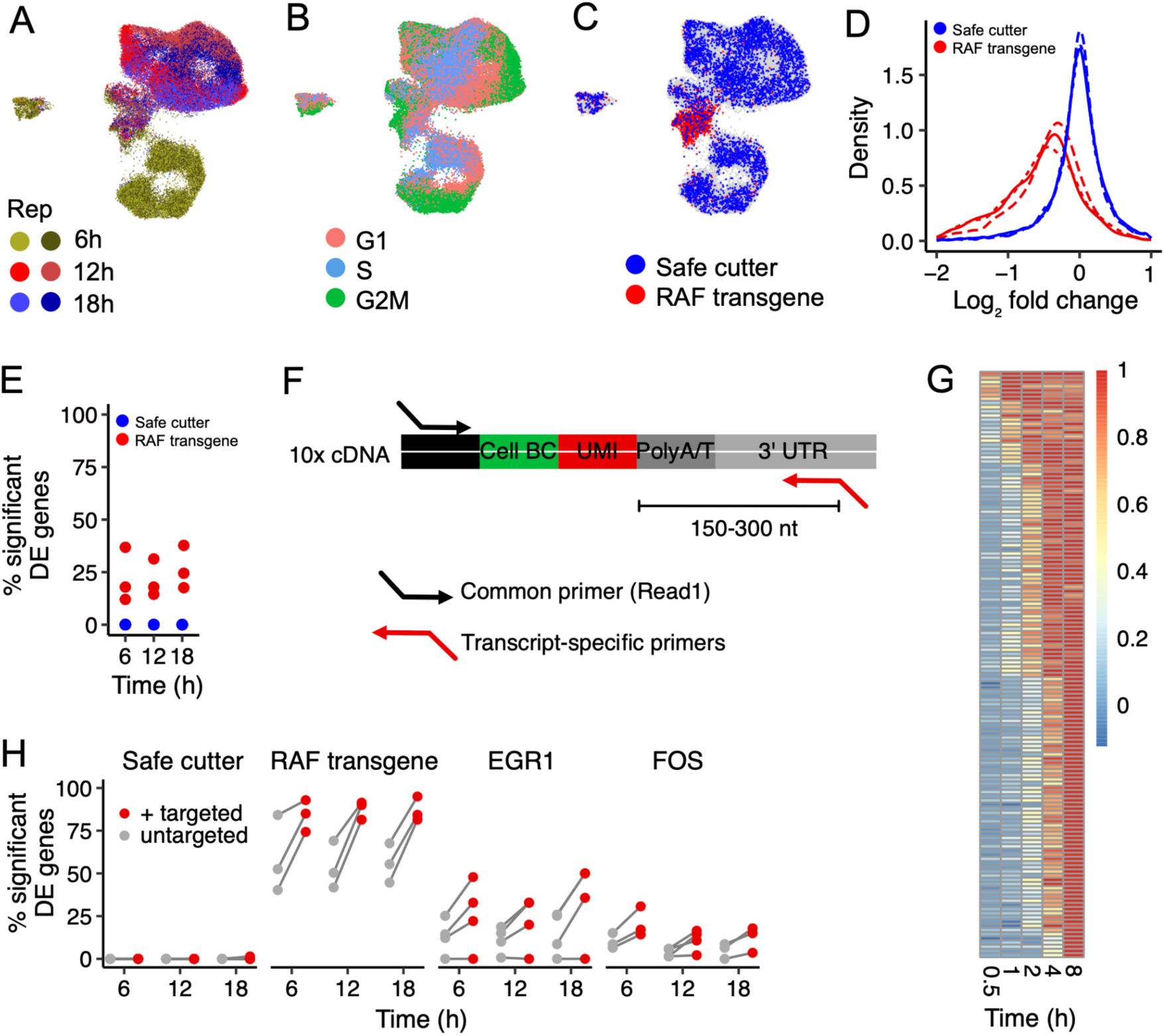
Targeted scRNA-Seq library amplification enhances Perturb-seq sensitivity. **(A)** UMAP integration of Perturb-seq samples from three different time points with two replicates each. **(B)** Distribution of cell cycle phases G1, S, and G2M on the integrated UMAPs. **(C)** Distribution of the safe cutter sgRNA cells and the RAF1-knockout cells on the UMAP clusters. **(D)** Histograms of the log_2_ fold change of differentially expressed genes between RAF1-knockout (red) and the safe cutter control cells (blue), relative to the non-target control cells. Dashed line: 6h time point, Dotted line: 12h time point, Solid Line: 18h time point. **(E)** Percentage of significantly differentially expressed genes in the RAF1-knockout cells from the total number of genes induced by RAF activation (adjusted p-value<0.05). Red = RAF1 sgRNAs, Blue = safe cutter sgRNAs. **(F)** Schematic of the modified TAP-seq approach. **(G)** Selection of 140 candidate genes for modified TAP-seq. Log_2_ gene expression fold changes from bulk RNA-Seq analysis of significantly induced genes after pulse induction of the RAF transgene for different times (FDR = 1%), normalized to gene-wise maximum Log_2_ fold changes. **(H)** Difference in percentage of significant differentially expressed genes from the 140 selected genes for the modified TAP-seq between targeted and untargeted approach after knockout of Raf transgene, EGR1, FOS, and safe cutter control.

### A modified TAP-seq approach enhances the sensitivity of Perturb-seq screens

Perturb-seq screens are becoming increasingly popular, but their analysis is subject to significant limitations, such as high sequencing costs, difficulty in detecting lowly expressed genes, and small effect sizes. Recently, a method called Targeted Perturb-seq or TAP-seq has been developed to overcome these challenges by PCR amplifying specific target genes of interest, directly from first-strand synthesized cDNA libraries (Schraivogel *et al*, 2020). Here, we developed an adaptation of TAP-seq which showed increased sensitivity without compromising the specificity of Perturb-seq screens. The modified TAP-seq approach uses the already PCR-amplified cDNA library as a template for a single step PCR target gene amplification in contrast to the original TAP-seq (Fig. 3F). This considerably reduces the risk of contamination of the original gene expression library through “off-priming” and makes it possible to retrospectively amplify additional target gene panels from the same gene expression library. The modified TAP-seq approach was used to amplify 140 target genes selected based on their activation and expression level following RAF1 induction (Fig. 3G). TAP-seq amplification did not alter the proportion of significantly deregulated genes in cells expressing safe cutter control sgRNAs, further confirming the specificity of the approach (Fig. 3H). For sgRNAs against the RAF1 positive control on the other hand, the proportion of significantly deregulated target genes increased substantially, demonstrating the increase in sensitivity gained by the modified TAP-seq approach. For most sgRNAs targeting the two example candidate TFs EGR1 and FOS, we observed a similar trend towards an increased proportion of significantly deregulated target genes (Fig. 3H).

### Number of target genes varies greatly between candidate TFs

After having established a modified TAP-seq protocol, we combined the targeted and untargeted Perturb-seq data to optimally analyze the 140 selected transcripts at high sequencing depth. In addition, it covers the entire transcriptome from the standard gene expression library. Figure 4 summarizes the results of a pseudo-bulk analysis of the combined targeted and untargeted Perturb-seq screens at 6h, 12h, and 18h after RAF1 induction, analyzed separately for each sgRNA. To ensure sufficient coverage, we excluded sgRNAs that were present in less than 30 cells. Consistent with the results shown in Figure 3H, the vast majority of the 10 safe cutter negative control sgRNAs yielded no significantly deregulated target genes over the three time points in comparison to non-target control sgRNAs. In contrast, sgRNAs targeting the positive control RAF1 yielded between 500 and 1,000 significantly deregulated target genes in all three time intervals post induction (Fig. 4A). Interestingly, the number of transcripts that were significantly deregulated after the perturbation of the 22 candidate TFs, varied greatly between the different TFs. While EGR1 perturbation for example, yielded hundreds of deregulated target genes, the perturbation of more than half of the candidate TFs did not lead to any significant deregulation of target genes above the cut-off.

**Figure 4.**
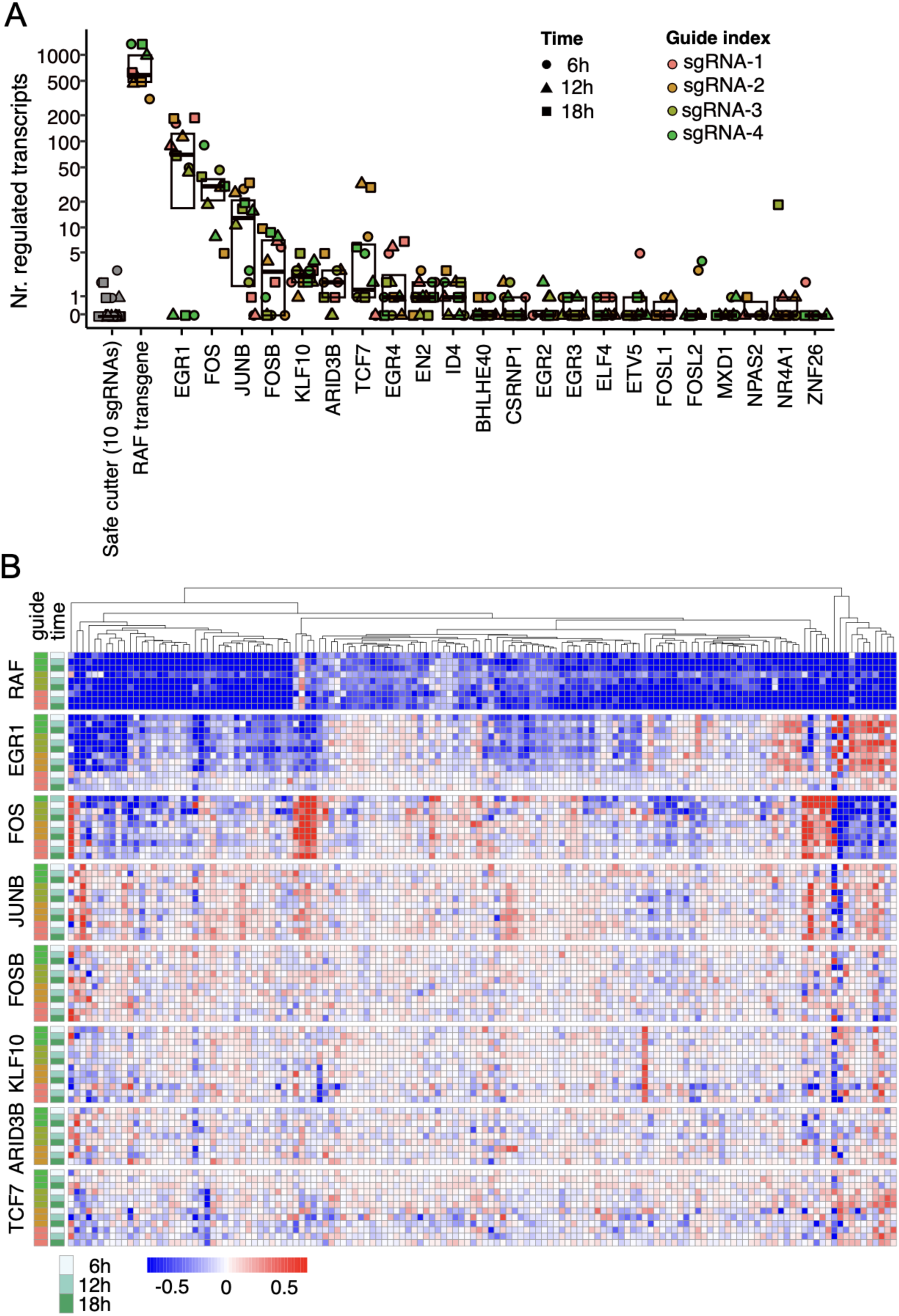
Summary of Perturb-seq screen results. **(A)** Number of significantly deregulated transcripts (adjusted p-value <0.05) following perturbation of the indicated candidate TF separated by sgRNAs and time points. **(B)** Heatmap of the log_2_ fold change of the 140 genes included in the modified TAP-seq for the perturbations with the strongest transcriptional response. The results from perturbed target genes separated by sgRNAs and time points are shown. Negative values indicate lower and positive values higher target transcript levels in the perturbed cells relative to cells expressing non-target control sgRNAs.

Figure 4B shows a heatmap of all candidate TFs whose perturbation resulted in significant deregulation of a subset of the 140 genes interrogated in the TAP-seq analysis. As expected, perturbation of the positive control RAF1 resulted in the inability of cells to activate RAF1 target genes upon 4OHT treatment, and hence in a negative log_2_ fold change of those genes relative to 4OHT-induced non-target control cells. The results also show that a large fraction of RAF1 targets are already deregulated after 6h of RAF1 induction in the RAF1- perturbed cells, while some are more strongly deregulated after 12h, consistent with the RAF1 target induction kinetics shown in Figure 1. We observed no major differences between 12 and 18h of RAF1 induction, suggesting that the full transcriptional response occurred after 12h (Schulze *et al*, 2001).

The transcriptional network downstream of RAF-MAPK is primarily co-regulated by 2 of the 22 investigated TFs, namely EGR1 and FOS. In contrast to RAF1, EGR1 and FOS perturbations mediating the highest number of deregulated target genes (Fig. 4A) appear to work through inhibition as well as activation of distinct target gene modules. Interestingly, EGR1 and FOS frequently seem to act orthogonally on overlapping sets of candidate genes, where one of the two TFs activates the expression of a target gene module, while the other TF inhibits it (Fig. 4B). How exactly do these two TFs regulate the RAF-MAPK transcription networks was further investigated below.

### De-novo construction of a TF core network identifies EGR1 and FOS as orthogonal regulators of the RAF-MAPK transcriptional response

Since the induction of target genes was similar over the different time points, we pooled all time points and sgRNAs against each individual TF, to define overall fold changes. Furthermore, we combined the significance of individual sgRNAs and time points by Fisher’s method to define significantly deregulated targets. Using this approach, we generated a “core heatmap” including only the candidate TFs (Fig. 5A). Examination of the differential expression of candidate TFs in response to self-regulation revealed a consistent, reasonably unexpected, down-regulation of most candidate genes. Since targeting Cas9 to early exonic regions introduces INDELs and thereby premature stop-gain mutations, the resulting transcripts are most likely degraded by nonsense-mediated decay, leading to the detected reduction in target mRNA levels (Kervestin & Jacobson, 2012). On the other hand, the observed up-regulation of the candidate TFs EGR1, FOS, and CSRNP1 could be explained by auto-regulatory feedback loops of those TFs in which the absence of the functional target protein leads to an up-regulation of its edited, stop-gain mutation containing mRNA. Consequently, an accurate assessment of perturbation efficiency should consider both elements, i.e. the downregulation of the perturbed target gene and an analysis of the differential expression of responding target genes.

**Figure 5.**
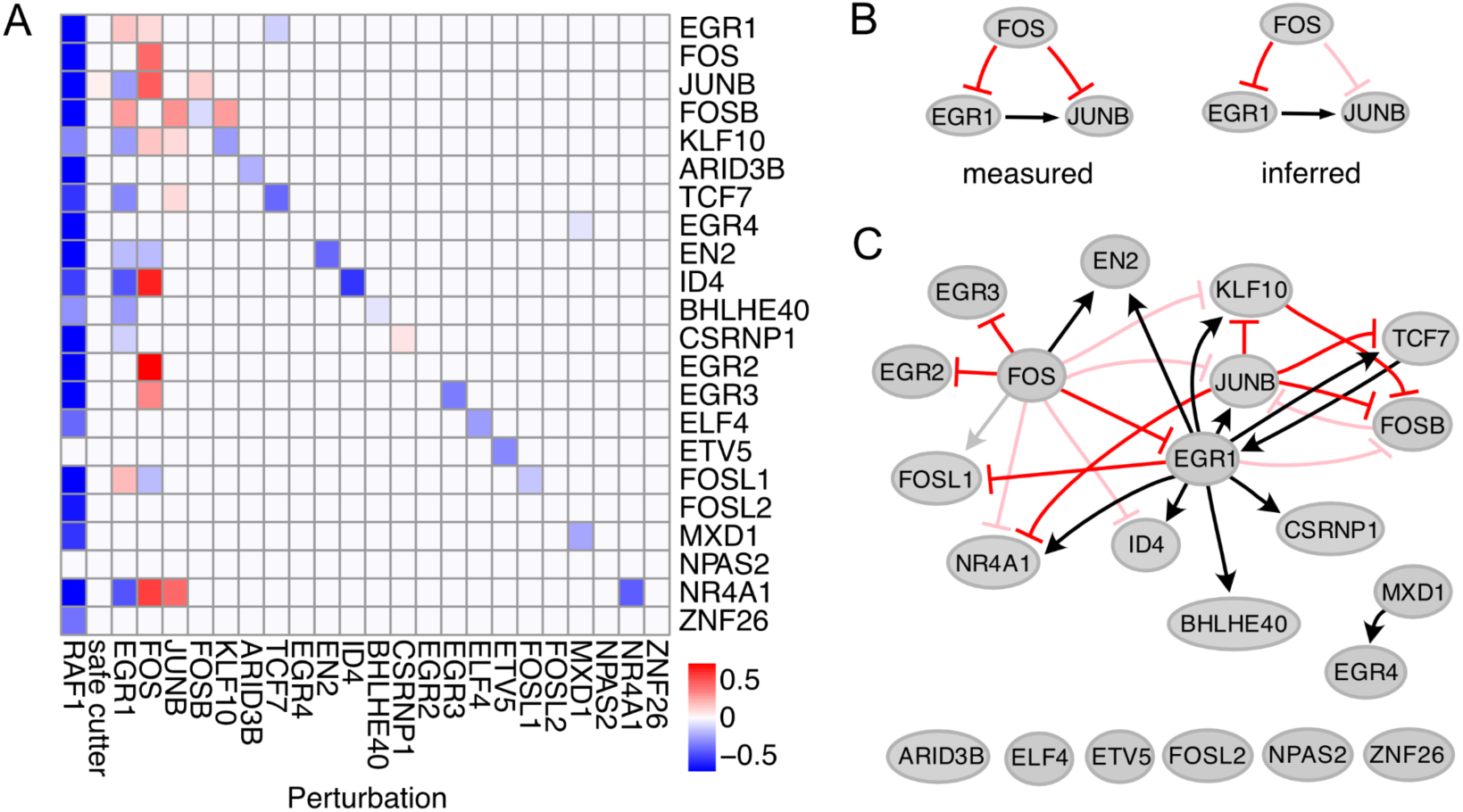
Model of transcriptional interactions between the 22 candidate TFs. **(A)** Heatmap of candidate TF expression changes following the perturbation of all 22 TFs and RAF1, showing the log_2_ fold change upon perturbation for the significantly differentially expressed TFs (adjusted p-value <0.05) **(B)** Example for the removal of edges from coherent feed forward loops: The measured perturbation data in A is compatible with a feed-forward loop, where FOS inhibits JUNB directly and indirectly, but it is also compatible with a cascade, where FOS inhibits JUNB via EGR1 only. To generate the most parsimonious network, we removed the feed forward loop from FOS to JUNB in the inferred network and termed it “indirect”. **(C)** De-novo model of the TF core network showing directional interactions between all perturbed TFs with inferred interaction type: Black = activating, Red = inhibiting. Transparent edges indicate the removed feed forward loops.

Consistent with earlier observations, EGR1 and FOS also act as central regulators in the TF core network, controlling the expression of several other candidate TFs in a predominantly orthogonal manner, where EGR1 activates the same candidate TF that is inhibited by FOS, this being the case for JUNB, KLF10, ID4 and NR4A1 (Fig. 5A). FOSL1 represents an example for the opposite co-regulation, where EGR1 inhibits its expression and FOS activates it. EN2 is the only candidate TF that is co-activated by EGR1 and FOS. Taken together, EGR1 was identified as a central activator within the TF core network, while FOS played a central inhibitory role, concurrently inhibiting EGR1 expression. This observation is consistent with the previously established roles of EGR1 and FOS as key regulators and emphasizes their central role in orchestrating the transcriptional response downstream of RAF1.

We derived a core transcriptional network from the perturbation data. Each TF represents a node that is connected to each significantly changed target by directional edges. We noted that the network contains several coherent feed forward loops, such as FOS inhibiting JUNB either directly or indirectly by inhibiting the upstream TF EGR1. From perturbation data alone, however, it cannot be deduced if the direct interaction, i.e. the feed forward loop, exists as both networks with and without feed forward loop result in the same perturbation response pattern in the heatmap (see example in Fig. 5B). Therefore, we chose the most parsimonious network, i.e. edges that correspond to the feed forward loops were removed (Fig. 5C).

### Directional interactions between TFs and their target genes

To check if the network is biologically plausible, we used time-series information from the bulk RNA-Seq experiments in Fig. 1. We expected that more upstream TFs are up-regulated earlier than their targets. Indeed, this is the case, with typical delays ranging from 15 min to 1h (Fig. 6A). However, the regulation of EGR1 by TCF7 represents a notable exception in that EGR1 is upregulated >5h before TCF7. Interestingly, EGR1 and TCF7 constitute the only feedback in the core network (Fig. 5C), which in turn leads to a non-identifiable network structure inherent to reconstruct qualitative networks from perturbation data. More precisely, target genes that respond to EGR1 and TCF7 perturbation, cannot be unequivocally assigned to one or both of the TFs, as all three resulting network topologies shown in Fig. 6B lead to exactly the same perturbation response pattern. However, because EGR1 was induced much earlier than TCF7, with a half-maximal induction time of 17 min versus 5h (Table 1), EGR1 was considered more “upstream” than TCF7, and therefore the feedback from TCF7 to EGR1 was removed. The final transcription factor network and the response times of each TF are shown in Figure 6C.

**Figure 6.**
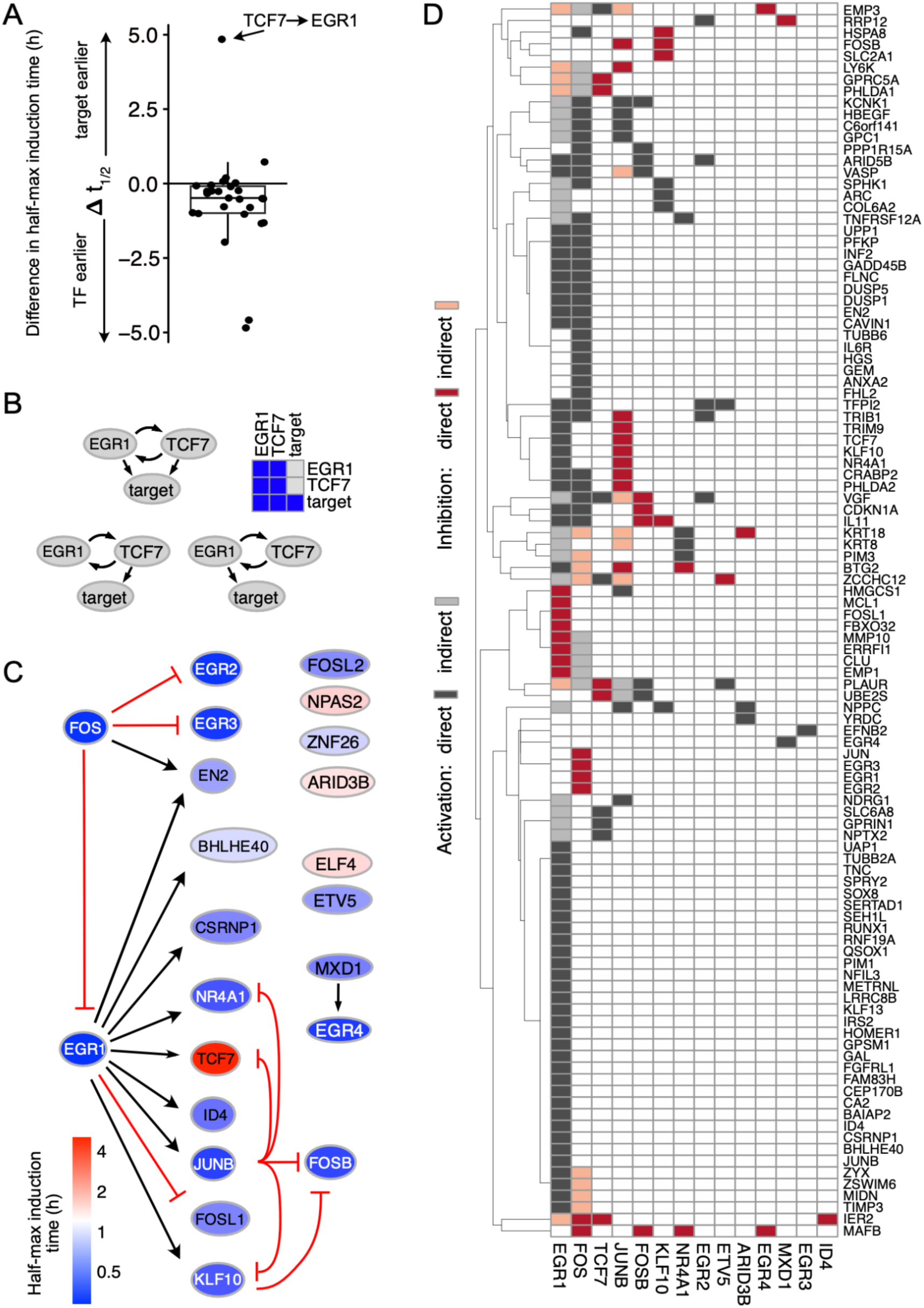
Identification of direct and indirect interactions from differentially expressed genes. **(A)** Difference in half-maximal induction times of TFs and their respective target genes in the core network was quantified. Positive values indicate that the target is induced first, negative values indicate that the TF is induced before the target. **(B)** The three displayed networks including the positive feedback of EGR1 and TCF7 and one target gene all result in the same stimulus-response pattern indicated in the heatmap. **(C)** De-novo model of interactions between the 22 candidate TFs with inferred interaction type: Black = activating. Red = inhibiting. Transcription factors are color-coded by their half-maximal induction time. **(D)** Summary of direct and indirect interactions between perturbed candidate TFs (x-axis) and their target genes (y-axis).

After the reconstruction of the TF core network, we analyzed the interaction of the network’s components with their target gene modules. For this, we added interaction to target genes whenever a TF perturbation leads to significant changes in expression of a target gene. We then repeated the network reconstruction procedure as above to remove feed forward loops, which resulted in the most parsimonious network. The removed feed forward loops were termed “indirect”. The resulting TF-target interaction map (Fig. 6D) clearly indicates that EGR1 and FOS are key mediators of the RAF-MAPK induced transcriptional response and that more downstream transcription factors regulate small, defined sets of target genes, either directly or indirectly.

### Bulk RNA-Seq analysis of single and combinatorial EGR1 and FOS knockout clones

The two central regulators of the transcriptional RAF1-MAPK response, EGR1 and FOS, were further investigated via bulk RNA-Seq of the responsive transcriptomes. For this purpose, we derived clonal EGR1 and FOS knockout lines from the parental HEK293ΔRAF1:ER cells and performed bulk RNA-Seq after RAF1 induction with 4OHT on each knockout clone. We found that the extent of EGR1 and FOS-mediated target gene regulation was largely concordantly covered by bulk RNA-Seq and Perturb-Seq analysis. Genes found down regulated in Perturb-seq were also downregulated in the bulk RNA-Seq analysis, and vice versa (Fig. 7A). Moreover, we observed a good quantitative correlation between the differential expression of the 140 transcripts quantified using the modified TAP-seq and bulk RNA-Seq following EGR1 or FOS knockout, respectively, with r-values ranging from 0.69 to 0.8 (Fig 7B). Taken together, these results confirm the high quality of the Perturb-seq datasets generated and used during the course of this project.

**Figure 7.**
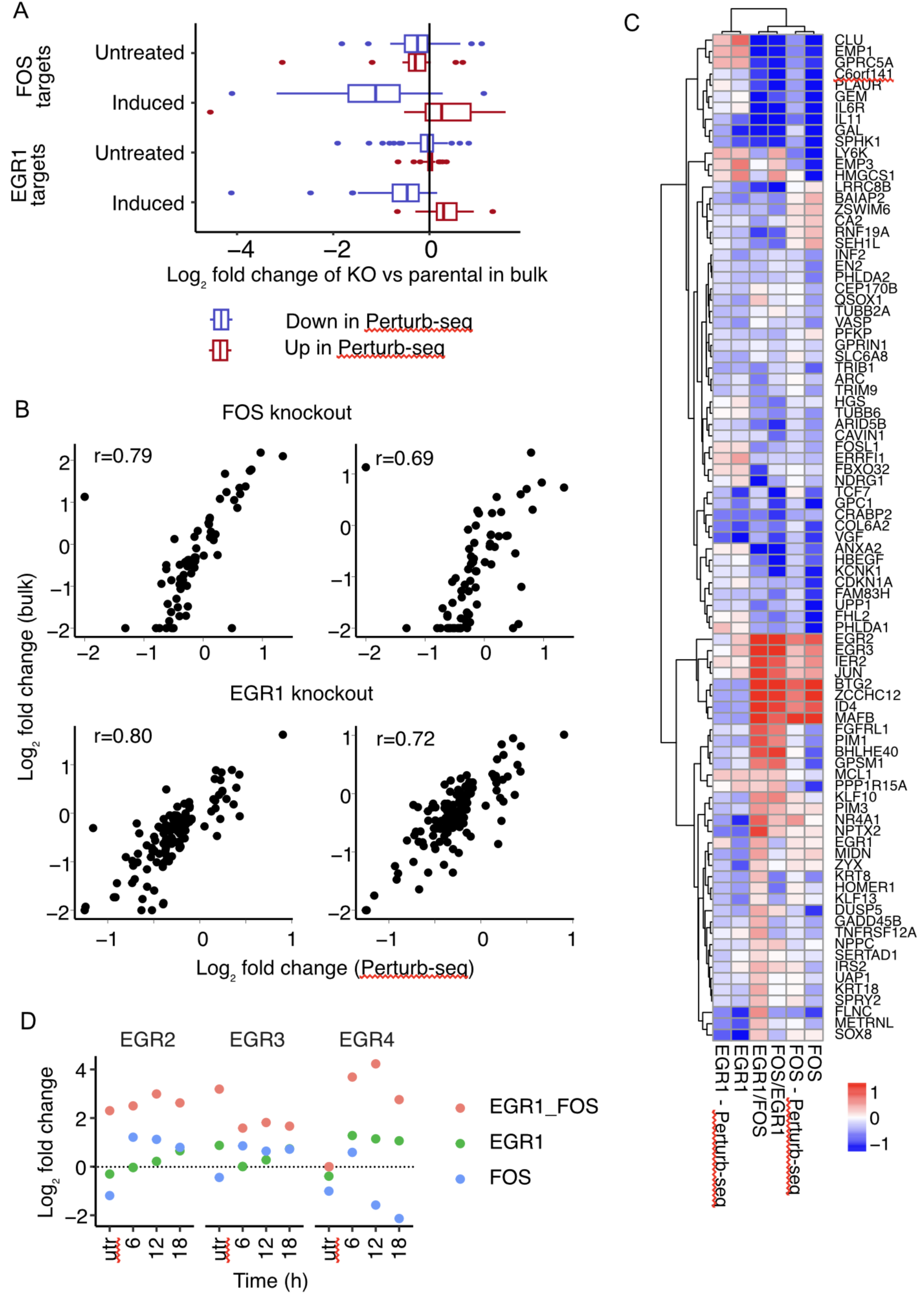
Bulk RNA-Seq analysis of individual and combinatorial EGR1 and FOS perturbations. **(A)** Average log_2_ fold changes for genes that are identified as up- and down-regulated target genes of EGR1 and FOS in perturb seq for bulk KO clones compared to parental clones. **(B)** Correlation between transcriptional changes detected via Perturb-seq and bulk RNA-Seq of EGR1- or FOS-perturbed cells from 2 different single knockout clonal lines of each gene. **(C)** Heatmap of transcriptional changes detected via bulk RNA-Seq from cells with individual or combinatorial EGR1 and FOS perturbations. Perturb-seq results from individual EGR1 and FOS perturbations are shown for comparison. **(D)** EGR2, EGR3, and EGR4 transcript levels determined via bulk RNA-Seq of EGR1/FOS single or double knockout clones, induced with 4OHT for 0h (utr), 6h, 12h, or 18h.

Starting from knockout clones of FOS and EGR1, we generated combinatorial knockout lines, in which both transcription factors were deleted in the same cell. The combinatorial knockout lines were used for bulk RNA-Seq after RAF1-induction, identical to the single knockout clones described above. The transcriptional changes detected via bulk RNA-Seq from EGR1 and FOS single and double knockout clones as well as the single knockout Perturb-seq results are shown in the heatmap in Fig. 7C. The good correlation between Perturb-seq and bulk RNA-Seq data from EGR1 and FOS knockout clones described above is also reflected in the heatmap, where Perturb-seq and bulk RNA-Seq expression profiles from single EGR1 and FOS knockouts clustered closely together. Interestingly, the combinatorial knockouts of EGR1 and FOS clustered closely together with the FOS only knockout, suggesting that globally the combinatorial knockout mimics the single FOS knockout. At a more granular level however, examples for the opposite scenario can be found as well in that the combinatorial knockout resembles the gene expression changes of EGR1 single knockout, exemplified by the LRRC8B, BAIAP2, ZSWIM6, CA2, RNF19A, SEH1L target gene cluster. We even found clusters in which the combinatorial expression pattern is contrary to both single knockout phenotypes, such as the FGFRL1, PIM1, BHLHE40, GPSM1 target gene cluster, in which the single knockout of EGR1 and FOS leads to a down-regulation while their combinatorial knockout leads to an up-regulation of those target genes (Fig. 7C).

EGR1 and FOS regulate overlapping sets of target genes, through highly target gene specific mechanisms. Analysis of the expression changes of EGR family members following single and combinatorial knockout of EGR1 and FOS revealed that FOS acts together with EGR1 to regulate EGR2, EGR3, and EGR4 in a synergistic manner (Fig. 7D). The expression levels of all three EGR family members increased only moderately after EGR1 single knockout but increased synergistically in the EGR1-FOS combinatorial knockout. Interestingly, the synergistic effects of EGR1-FOS double knockouts are most striking for EGR2 and EGR3 in non-induced cells, while we observed no effect on EGR4. On the other hand, FOS single knockout even leads to EGR4 down-regulation and EGR1 knockout to a modest EGR4 up-regulation in RAF1-induced cells and their combinatorial knockout leads to a strong synergistic up-regulation of EGR4.

## Discussion

The MAPK signaling pathway is a key cellular signaling cascade that has a significant impact on many human diseases (Kim & Choi, 2010). Extensive research has focused on characterizing the upstream regulators of this pathway, which has led to a detailed understanding of the network (Oda *et al*, 2005). This research has uncovered the mechanisms by which the signaling pathway generates distinct temporal patterns of activity that lead to different cell fate decisions (Ram *et al*, 2023a; Blum *et al*, 2019; von Kriegsheim *et al*, 2009; Vaudry *et al*, 2002). However, the mechanisms by which these activity patterns are decoded and translated into different transcriptional responses remain poorly understood (Ram *et al*, 2023b). Previous investigations have identified small crucial components of the decoding of long lasting stimuli. These include feed-forward loops, for example in the post-translational stabilization of FOS by ERK (Murphy *et al*, 2002), and long mRNA half-lives of some early genes that filter short pulses (Uhlitz *et al*, 2017). Nevertheless, our understanding of how the transcriptional response downstream of RAF-MAPK is organized by the transcription factor network is limited. Transcriptional network modeling based on genetic perturbation data offers a powerful approach to address this shortcoming (Stelniec-Klotz *et al*, 2012). Targeted Perturb-seq now provides a tool for measuring transcriptional responses to genetic perturbations on a large scale and provides the data sets required for de novo construction and refinement of transcriptional network models to decipher the transcriptional networks associated with the RAF-MAPK signaling pathway.

In this study, we employed a topology-based modeling approach with data from Perturb-seq experiments. Using “Occam’s razor” principle, the simplest network configuration that aligned with the measured data was created. Specifically, we eliminated coherent feed-forward loops, where a TF might regulate a target gene both directly and via another TF. A similar technique is also used in reverse engineering of transcriptional networks from correlation with ARACNE, which removes the least supported edge in a triangle (Margolin *et al*, 2006). While coherent feed-forward loops are known to exist in transcriptional networks and may possess biologically significant functions (Milo *et al*, 2002), their removal aided in deriving a more streamlined network structure. To gain a clearer understanding of the biological functions of feed-forward structures, orthogonal CRISPR approaches, involving the knockout of one gene combined with the activation of a second gene in the same cell (Boettcher *et al*, 2018), could be employed in future investigations.

The extensive Perturb-seq datasets generated in this study facilitated the construction of a “core network”, outlining directional interactions among the 22 perturbed candidate TFs. By eliminating feed-forward loops within this core network, as well as in the extended network comprising all significantly deregulated target genes of the perturbed TFs, we could distinguish likely direct from indirect interactions. This process allowed the derivation of a more parsimonious network structure. Integration of these minimal interaction networks with TF expression kinetics further enabled the removal of positive feedback loops. Notably, the immediate early genes EGR1 and FOS emerged as the most interactive upstream co-regulators, with orthogonal regulatory influence on the expression of other TFs and downstream targets within the network.

One important function of several delayed early and secondary response genes is negative feedback regulation (Avraham & Yarden, 2011; Legewie *et al*, 2008). We noticed that several feedback regulators such as dual-specificity phosphatases DUSP1 and DUSP5 are dependent on EGR1 and FOS (Fig. 6D and 7C). Therefore, loss of EGR1 and FOS may limit feedback regulation and lead to increased MAPK signaling. This may also be reflected in the inferred gene regulatory network, in which we observe an inhibitory influence of FOS and EGR1 on multiple immediate early transcription factors and immediate early genes. To further decipher the complex feedback regulation between the transcription network and signal transduction, perturbation datasets that additionally measure the activity of signaling are required (Dessauges *et al*, 2022; Genolet *et al*, 2022).

The framework presented here serves as a basis for the systematic study of transcriptional network dynamics. Its scalability and effectiveness will enable future research into transcriptional networks beyond the specific context of RAF-MAPK signaling, thereby allowing the disentanglement of the regulatory networks associated with all types of cellular processes and diseases.

## Methods

### Vector Maps

All in one Cas9 vector with 10x Genomics capture sequence 1, 5′-GCTTTAAGGCCGGTCCTAGCAA-3′, in the stem-loop of the Cas9-tracr sequence (pMB1-10x) was used (Replogle *et al*, 2020). The plasmid map is provided in GenBank format (Supplementary Data 1).

### HEK293ΔRAF1:ER cell culture

HEK293ΔRAF1:ER cells (Cagnol *et al*, 2006) containing a tamoxifen inducible fusion of the kinase domain of RAF1 (Samuels *et al*, 1993; McMahon, 2001) reviewed in (Samuels *et al*, 1993; McMahon, 2001) were cultured in complete DMEM low glucose without phenol red supplemented with 10% fetal bovine serum (Pan Biotech) and 1% antibiotics (pen/strep). Lenti-X 293T cells (Takara) were cultured in complete DMEM supplemented with 10% fetal bovine serum and 1% antibiotics (pen/strep).

### Bulk RNA-sequencing data generation and preprocessing

Total RNA was extracted with TRIzol. Sequencing libraries were prepared using Illumina TruSeq mRNA Library Prep Kit v2 and sequenced on Illumina HiSeq 2000. Raw reads were processed using the snakemake-workflows rna-seq-star-deseq2 pipeline, v1.2.0, and counts were subsequently analyzed using DESeq2 in R.

### Cas9 library design

Target genes were selected based on RNAseq data after induction of HEK293ΔRAF1:ER cells with 0.5 µM 4-hydroxytamoxifen (4OHT) at different time points (Fig. 1). The sgRNA library consisted of 4 sgRNAs per gene with 10 non-target control sgRNA and 10 safe cutters sgRNAs. The sgRNA sequences were selected from the Brunello genome wide library (Doench *et al*, 2016). 4 positive-control sgRNAs against the Raf-transgene were designed using CRISPick (Doench *et al*, 2016; Sanson *et al*, 2018) (Supplementary Table 1).

### Cas9 library cloning

The selected 20-nt target specific sgRNA sequences were cloned into the pMB1-10x library vector (Supplementary Data 1) by Gibson Assembly (Gibson *et al*, 2009). sgRNA template sequences of the format: 5′-GGAGAACCACCTTGTTGG-(N)20-GTTTAAGAGCTAAGCTGGAAAC-3′ were synthesized in a pooled format on microarray surfaces (GenScript Biotech, Inc.). Oligo pools were PCR-amplified using Phusion Flash High-Fidelity PCR Master Mix (ThermoFisher Scientific) according to the manufacturers protocol with 1 ng/μL sgRNA template DNA, 1 μM forward primer (5′-GGAGAACCACCTTGTTGG-3′), and 1 μM reverse primer (5′-GTTTCCAGCTTAGCTCTTAAAC-3′) in 50 µL total volume. The following cycle numbers were used: 1× (98 °C for 3 min), 16× (98 °C for 1 s, 54 °C for 15 s, 72 °C for 20 s) and 1× (72 °C for 5 min). PCR products were purified using NucleoSpin columns (Macherey-Nagel). The library vector pMB1-10x was prepared by restriction digestion with AarI (Thermo Fisher) at 37 °C overnight. The digestion reaction was run on a 1% agarose gel followed by excision of the digested band and purification via NucleoSpin columns (Macherey-Nagel). 100 ng digested pMB1-10x and 2.4 ng amplified sgRNA library insert were assembled using Gibson Assembly Master Mix (NEB) in a 20 μL reaction for 30 min. The reaction was purified using P-30 buffer exchange columns (BioRad) that were equilibrated 5x with H_2_O and the eluted volume was transformed into 20 µL of MegaX DH10β cells (Thermo Fisher) by electroporation. *Escherichia coli* were recovered and cultured overnight in 100 mL LB medium with 100 μg/mL ampicillin. The plasmid library was extracted using Midiprep (Qiagen). In parallel, a fraction of the transformation reaction was plated and used to determine the total number of transformed clones. The coverage was determined to be 1,643x clones per sgRNA ensuring even representation of all library sgRNA sequences and their narrow distribution (Fig. 2B). The quality of the cloned sgRNA library was determined by NGS on an Illumina Miniseq (See below). MAGeCK (Li *et al*, 2014) was used for library alignment. Narrow distribution of sgRNA sequences was confirmed with read counts for 96% of sgRNA sequences falling within a single order of magnitude.

### Lentivirus production

Lenti-X 293T cells (Takara) were seeded at 65,000 cells per cm^2^ in 10 mL media (DMEM, 10% FBS, 1% pen/strep) in a 10 cm dish and incubated overnight at 37 °C, 5% CO_2_. On the next day, 5 μg sgRNA library plasmid, 2 μg psPAX2 (Addgene #12260), 2 μg pMD2.G (Addgene #12259), and 36 μL Turbofect (Thermo Fisher) were mixed into 1.8 mL serum-free DMEM (Gibco), vortexed briefly, incubated for 20 min at RT, and added to the cells. At 48 and 72h post-transfection, the supernatant was harvested, passed through 0.45 um filters (Millipore), and 20x concentrated using LentiX Concentrator (Takara) according to the manufacturer’s instructions. Aliquots were stored at −20 °C.

### Direct capture Perturb-seq CRISPR screens

HEK293ΔRAF1:ER cells were transduced with lentivirally packaged sgRNA library at a multiplicity of infection (MOI) = 0.2 and 1,000x coverage in 2 replicates. The low MOI and high coverage were used to reduce the frequency of multiple-infected cells thus only one gene was knocked out in each cell and ensure even distribution of the sgRNA library. Cells were then cultured in DMEM low glucose without phenol red with 10% FBS (Pan Biotech) and 1% pen/strep (Sigma-Aldrich) in a 37 °C incubator with 5% CO_2_. 48h after transduction, transduced cells were selected with puromycin (2 μg/mL) for 96h. After selection, the top 20% mcherry positive cells were sorted 8 days post infection using a BD FACSAria II flow cytometer to increase sgRNA capture efficiency by the 10x Genomics Gel Beads. A total of 3 million cells were sorted. For the Perturb-Seq screen, 500,000 of the sorted cells were reseeded in full medium in 3 wells of a 12 well plate and incubated at 37 °C, 5% CO_2_. At day 10 post infection, the sorted cells were stimulated with 0.5 µM 4OHT (Sigma-Aldrich, H7904) for 6, 12, and 18h. After the incubation time, the cells were harvested followed by scRNA-Seq following the 10x Genomics Chromium Next GEM Single Cell 3’ Reagent Kits v3.1 (Dual Index) with Feature Barcode technology for CRISPR Screening protocol.

### Pooled proliferation CRISPR screen

The remaining 2.5 million sorted HEK293ΔRAF1:ER cells from day 8 post infection were reseeded in full medium in a 6 well plate and incubated at 37 °C, 5% CO_2_. On day 18 post infection, 0.5 µM 4OHT was added for 48h to induce the RAF1 transgene expression and apoptosis of the induced cells. After 48h, the dead cells were detached and removed with the medium. The living cells were reseeded in tamoxifen containing medium and incubated at 37 °C, 5% CO_2_ for 48h more to increase the cell selection efficiency. Aliquots of 500,000 cells from the 48h induction time point were taken. The cells were centrifuged, and the cell pellets were frozen down for later analysis via NGS.

### Genomic DNA extraction and PCR recovery of gRNA sequences

The genomic DNA (gDNA) was extracted from the HEK293ΔRAF1:ER cells using the Qiagen Genomic DNA extraction kit according to the manufacturer’s instructions. Two nested PCR reactions were performed to amplify the sgRNA cassette from the extracted gDNA. For the first PCR reactions, 5 μg gDNA, 0.3 μM forward (5′-GGCTTGGATTTCTATAACTTCGTATAGCA-3) and reverse (5′-CGGGGACTGTGGGCGATGTG-3′) primer, 200 μM dNTP mix, 1x Titanium Taq buffer and 2 μL Titanium Taq polymerase (Takara) were mixed in 50 µL total volume. The PCR reaction was run using the following cycles: 1x (94 °C, 3 min), 20x (94 °C, 30 s, 65 °C, 10 s, 72 °C, 20 s), 1x (68 °C, 2 min). For the second PCR reactions, 5 μL first-round PCR, 0.5 μM forward (5′-AATGATACGGCGACCACCGAGATCTACACACACTCTTTCCCTACACGACGCTCTTCCGATCTTCCCTTGGAGAACCACCTTGTTGG-3′) and reverse (5′-CAAGCAGAAGACGGCATACGAGAT-(N)_6_-GTGACTGGAGTTCAGACGTGTGCTCTTCCGATC-3′) primer where (N)_6_ is a 6 nt index for sequencing on the Illumina Miniseq platform, 200 μM dNTP mix, 1x Titanium Taq buffer and 2 μL Titanium Taq (Takara). PCR cycles were: 1x (94 °C, 3 min), 20x (94 °C, 30 s, 55 °C, 10 s, 72 °C, 20 s), 1x (68 °C, 2 min). The PCR product (325 bp) was purified from a 1% agarose gel via NucleoSpin columns (Macherey-Nagel). NGS was performed on an Illumina Miniseq using a MiniSeq Mid Output Kit (300-cycles) using paired-end 150 strategy according to the manufacturer’s instructions.

### Proliferation screen data analysis

The proliferation screen data analysis was performed using MAGeCK (Li *et al*, 2014). In short, sgRNA read count files were computed from the raw CRISPR fastq files using the count function. The MAGeCK MLE command was then used to calculate the MAGeCK Beta score, Wald-P values, and false discovery rates for enrichment and depletion of each guide at day 20 and day 22 after tamoxifen induction compared to the plasmid library. Wald-P values were adjusted using Bonferroni correction method in R (Supplementary Table 2).

### scRNA-Seq screen data analysis

Cell Ranger (10x Genomics) Version 6.1.1 was used for scRNA-Seq data processing (https://support.10xgenomics.com/single-cell-gene-expression/software/pipelines/latest/using/count). Sequencing reads coming from the gene expression library were mapped to the GRCh38-1.2.0 genome reference compiled by 10x Genomics for Cell Ranger. Guide RNA reads were mapped simultaneously to a sgRNA feature reference. The combination of standard and targeted RNA-Seq was processed by pooling the fastq files and subsequent Cell Ranger analysis, thereby avoiding duplicate counts for the same molecules as reads with the same UMIs are collapsed. Count matrices were then used as input into the Seurat R package (Hao *et al*, 2021) to perform downstream analyses. Differential expression was called based on pseudo bulks using the R-library glmGamPoi version 1.10.2 (Ahlmann-Eltze & Huber, 2021).

### Gene expression library amplification (TAP-seq)

For the generation of suitable primers, we used the BAM file of an untargeted 10x run (6h 4OHT) in the TAP-seq R package (https://github.com/argschwind/TAPseq) and workflow that is delineated in the package vignette (Schraivogel *et al*, 2020). Deviations from the TAP-seq workflow were as follows: (i) We only used the inner primer generation procedure, i.e. at 150-300bp from inferred poly(A) sites. (ii) Originally primers for one major poly(A) site per gene were generated, instead we generated primers for all top poly(A) sites amounting to >70% of the total poly(A) score (i.e. coverage) per gene. (iii) Additional filter steps on poly(A) level, i.e., removal of minor poly(A) sites in adjacent poly(A) signals within 100bp, and on primer level, i.e., removal of redundant primers within 500bp of poly(A) signal and manual filter of badly designed primer e.g., in non-expressed regions. The final primer list is provided in the Supplementary Table 3. Targeted primers were ordered from IDT in desalted format with 5’-GTGACTGGAGTTCAGACGTGTGCTCTTCCGATCT-3’ as PCR handle at 5’ ends. 100 µL PCR was performed as follows: 100 ng amplified cDNA (Step 2.3 of 10x Genomics 3’scRNA), 2.5 µL 100 µM Pooled targeted Primer Mix, 4 µL 10 µM partial Read 1(sequence from 10x Genomics manual, ordered as normal (desalted) primer from IDT 5’-CTACACGACGCTCTTCCGATCT-3’, 50 µL KAPA HiFi HS RM (Roche, KK2601). The PCR was performed with the following cycle numbers: 1x (95 °C, 3 min), 10x (98 °C, 20 s, 67 °C, 60 s, 72 °C, 60 s), 1x (72 °C, 5 min). The PCR product was cleaned using SPRIselect beads (Beckmann Coulter) as a double-sided size selection (as described in the Tips & Best Practices Section of a typical 10x Genomics procedure) with 0.6x as 1^st^ SPRI and 1.2x as 2^nd^ SPRIselect steps. For adding indices, maximally 10 ng of the first cleaned PCR product were then mixed with 20 µL dual Index TT Set A from 10x genomics, 50 µL KAPA HiFi HS RM (Roche, KK2601) in a 100 µL reaction. The PCR was performed with the following cycle numbers: 1x (95 °C, 3 min), 1x (98 °C, 45 s), 10x (98 °C, 20 s, 54 °C, 30 s, 72 °C, 20 s), 1x (72 °C, 1 min). As the amplicons are bigger now, we changed the SPRI concentration to 0.55x as 1^st^ SPRI and 1.2x as 2^nd^ SPRIselect steps. Library quality was assessed using the TAPE station.

### EGR1 and FOS knockout clonal line production

HEK293ΔRAF1:ER single knockout clones for FOS and EGR1 genes were generated using CRISPR-Cas9. Two sgRNAs per gene were designed to target 500 bp sequences surrounding the 5’ end using CRISPOR (Concordet & Haeussler, 2018). The sgRNAs and their reverse complements were synthesized and cloned into the px459 vector (Ran *et al*, 2013). For the EGR1 gene knockout, the following oligo sequences were used: 5’-CACCGGGCCATGTACGTCACGACGG-3’ and 5’-CACCGGGACAACTACCCTAAGCTGG-3’ targeting the promoter and the exon regions respectively. For the FOS gene knockout, the following oligos sequence was used to target the promoter region; 5’-CACCGGATTAGGACACGCGCCAAGG-3’ and the exon region, 5’-CACCGGAGAGAGGCTATCCCCGGCCG-3’. The oligo for FOS exon contains an added G at the 5’ end of the gRNA to facilitate U6 promotor mediated transcription. Cells were transfected with these vectors following the Lipofectamine 2000 (ThermoFisher Scientific) and selected with puromycin (250 ng/mL) for 36h starting 24h after transfection. Following selection, we performed clonal dilution. Cells were seeded in 96 well plates and wells with individual clones were screened via PCR to identify successful 5’ end deletions. Positive hits were further validated at RNA and protein levels using qPCR and Western blot, to confirm the absence of FOS and EGR1 gene expression.

Starting from single knockout clones of EGR1 and FOS, double knockout clones were generated using Alt-R™ S.p. Cas9 Nuclease V3 from IDT used with guides designed with the manufacturer’s design tool (https://eu.idtdna.com/site/order/designtool/index/CRISPR_CUSTOM) (Supplementary Table 4). Transfection was performed following the Lipofectamine CRISPRMAX (ThermoFisher Scientific), using less RNA amounts depending on the number of sgRNAs used per transfection. Two days after transfection, clonal dilution was performed. Isolation of gDNA was done with Quick-DNA-96 Kits (Zymo), for PCRs KAPA HiFi HS RM (Roche) and different primer sets spanning regions out and/or inside expected deletions were used, PCR product size was analyzed with TAPE Station from Agilent. Clones with successful deletions based on the PCR result were then analyzed using Western blots with antibodies against FOS or EGR1 and pERK antibody (Cell Signaling Technology) to identify clones that lack expression of the respective TFs and still induce MAPK signaling when the RAF1-CR3 kinase domain is activated with 4OHT. KO clones were cultivated in low glucose DMEM without Phenol red (Sigma-Aldrich) supplemented with stable Glutamine (PAN-Biotech) and FBS (PAN-Biotech).

### Bulk RNA-Seq of single and combinatorial EGR1 and FOS knockout clonal lines

For bulk RNA sequencing cells were treated with 0.5 µM 4OHT for 6h, 12h, and 18h. To account for differences in cell density, two solvent control wells were collected each at first and last lysing time of the treatments. RNA isolation was done with RNeasy Kits from Qiagen without any DNA elimination. RNA concentration was measured with the Implen nanophotometer and 250 to 450 ng per sample were used for Library preparation with QuantSeq 3’ mRNA-Seq V2 (Lexogen). For Index PCR, 18 cycles were used according to the manual. Upon quantification with Universal KAPA Library Quantification Kit for Illumina (Roche) Libraries were pooled and sent for sequencing.

### Processing of bulk, multiplexed QuantSeq data

bcl2fastq (v2.20.0 by Illumina) was used to demultiplex and convert raw sequencing data to fastq files. We designed a Snakemake (v7.18.2) workflow in which BBMap’s BBDuk (v39.01) was used to trim adapters, STAR (v2.7.10b) to align reads to the GENCODE GRCh38.p13 (v39) geneset, umitools (v1.1.4) to extract and deduplicate UMIs, and subread’s featureCounts (v2.0.6) to count mapped reads on gene level.

## Data availability

Raw and processed transcriptome data is available at GEO under the accession number <u>GSE250559</u>. Data processing scripts and raw input data for the data processing scripts are available at Zenodo at the following https://zenodo.org/records/10493550.

## Author contributions

**Ghanem El Kassem**: Optimized and executed Perturb-seq and proliferation screens, cloned sgRNA library, writing

**Anja Sieber**: Designed and conducted targeted Perturb-seq experiments, transcriptome experiments, and generated and analyzed double KO lines

**Bertram Klinger**: Designed targeted Perturb-seq experiment

**Florian Uhlitz**: Performed time-series and pulsed transcriptome experiments

**David Steinbrecht**: Performed transcriptome analysis

**Mirjam van Bentum**: Generated and analyzed KO lines

**Jasmine Hillmer**: Performed data analysis

**Jennifer von Schlichting**: Performed data analysis

**Reinhold Schäfer:** Characterized cell lines and clones, manuscript revision

**Nils Blüthgen**: Supervision, funding acquisition, conceived the study, data analysis, writing

**Michael Boettcher**: Supervision, designed sgRNA library, designed experiments, funding acquisition, writing

## Disclosure and competing interest statement

The authors declare that they have no conflict of interest.

## Acknowledgments

This work was supported by the European Social Fund (ZS/2016/08/80642) to MB, by the Einstein Foundation Berlin, grant EVF-BIH-2019-512 to NB, by the Deutsche Forschungsgemeinschaft (DFG), grants TRR186/A18 and RTG2424 to NB, by the Deutsche Krebshilfe (DKH), grant 70114307 to BK and RS and by the German Cancer Consortium (DKTK) Young Investigator grant to BK.

## Supplementary material

Supplemantary Data 1: Map of pMB1-10x vector for CRISPR screen sgRNA expression.

Supplementary Table 1: sgRNA Library sgRNA sequences

Supplementary Table 2: Proliferation screen read count table and MAGeCK MLE values

Supplementary Table 3: Modified TAP-seq targets inner primers for gene expression library amplification

Supplementary Table 4: sgRNA sequences used for the generation of EGR1 and FOS double knockout clonal lines

## Notes

https://www.ncbi.nlm.nih.gov/geo/query/acc.cgi?acc=GSE250559

https://zenodo.org/records/10493550

### Competing Interest Statement

The authors have declared no competing interest.

