## Supplementary material for "Targeted Perturb-seq Reveals EGR1 and FOS as Key Regulators of the Transcriptional RAF-MAPK Response": Fig. S

### Supplementary figures

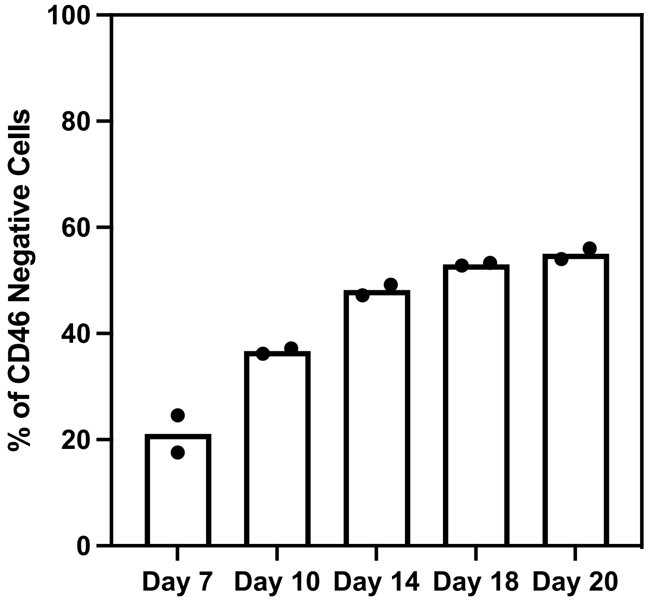


**Supplementary Figure 1 - CD46 knockout kinetics in HEK293ΔRAF1:ER cells at different time points after lentiviral infection.** Percentage of CD46 knockout cells was determined via flow cytometry analysis of >10,000 cells stained with CD46 antibodies (Miltenyi).

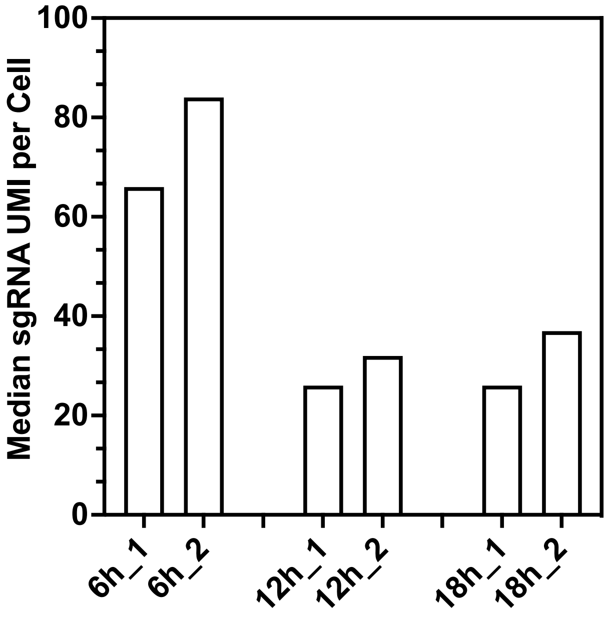


Supplementary Figure 2 - Median sgRNA UMI counts per cell detected in the respective Perturb-seq samples.

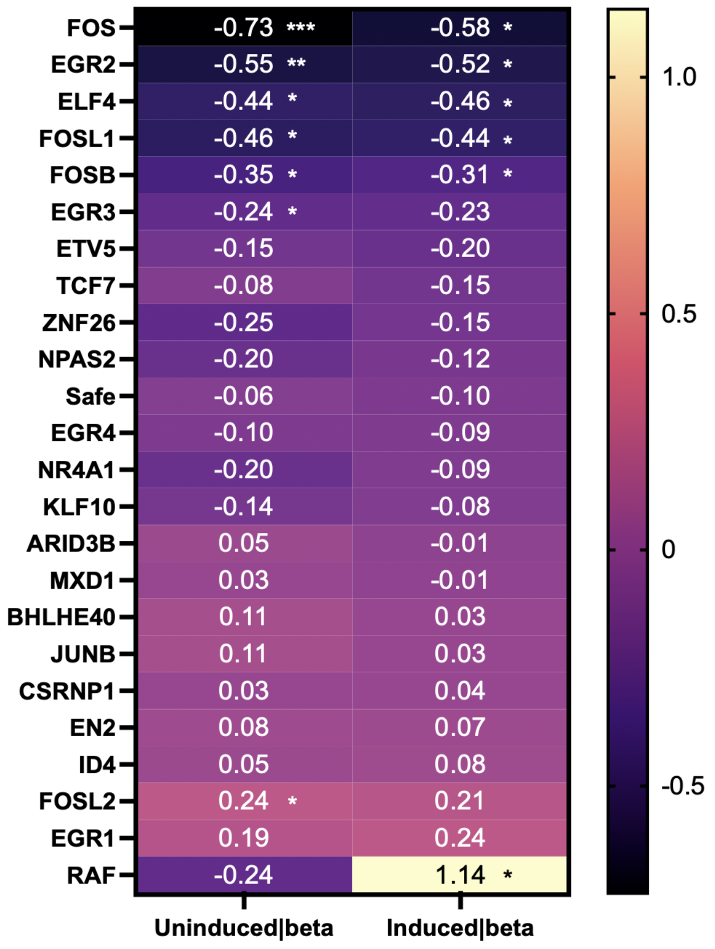


**Supplementary Figure 3 - CRISPR/Cas9 proliferation screen in HEK293ΔRAF1:ER cells.** MAGeCK MLE beta scores and corresponding significance are shown. * wald FDR <0.05, ** wald FDR <0.01, *** wald FDR <0.001
